# Alterations to the yeast chromatin landscape during hypoxia

**DOI:** 10.64898/2026.09.13.751330

**Authors:** Winny Sun, Maraki Negesse, Chase Andre, Virginia Murray, Josephine J. Szczyrbak, Erin M. Green

**Affiliations:** Department of Biological Sciences, University of Maryland Baltimore County, Baltimore, MD, 21250; Marlene and Stewart Greenebaum Comprehensive Cancer Center, University of Maryland School of Medicine, Baltimore, MD 21201

**Keywords:** chromatin, histone modifications, acetylation, methylation, hypoxia, yeast

## Abstract

When cells encounter low oxygen conditions, signaling events target key transcription factors to reprogram gene expression to promote cell survival and adaptation under this challenging environmental stress. The substantial metabolic changes that occur in hypoxia also drive changes to the chromatin environment due to altered cofactor availability and direct oxygen sensitivity of chromatin-modifying enzymes, among other factors. Given the links between hypoxic cellular environments and numerous pathophysiological processes, including tumor progression and metastasis, cardiovascular disorders, and aberrant development, it is critical to understand the chromatin states and chromatin-based regulatory mechanisms associated with cellular responses to hypoxia. Here, we used the model eukaryote *Saccharomyces cerevisiae* to interrogate the requirement for key transcription factors and chromatin regulators in survival during hypoxia and we assessed the distribution of major histone modifications associated with the response to stress, H3K9 acetylation and H3K4 methylation, throughout the genome during hypoxia. Our results show that only a small number of chromatin modifiers in yeast contribute to survival in hypoxia; however, there are substantial changes to the abundance and distribution of both H3K9ac and H3K4me3 during hypoxic growth. Specifically, H3K9ac levels are reduced throughout the genome though still maintain association with transcriptionally active genes. H3K4me3 has increased abundance genome-wide and shows a wider distribution with greater abundance in coding sequences particularly of genes upregulated in hypoxic conditions. Altogether, our data demonstrate substantial changes to the chromatin landscape in hypoxic conditions in the budding yeast model, providing key insights into physiological chromatin changes caused by limiting oxygen in the environment.

## INTRODUCTION

Molecular pathways that regulate gene expression in response to the environment often signal to transcription factors and chromatin regulators, including histone modifiers such as methyltransferases and acetyltransferases (1–3), to direct gene expression and promote survival or differentiation under the new conditions. Hypoxia, or low oxygen conditions, is frequently encountered by cells and results in massive metabolic and genomic reprogramming in a manner dependent on oxygen concentration and duration of exposure (4,5). Changes to the post-translational modification (PTM) of histones including methylation and acetylation, among others, reflect the altered metabolic state of the cells upon hypoxia exposure and are associated with hypoxia responsive gene expression programs that support survival and adaptation to low oxygen (6–8). In mammalian cells, the reprogrammed chromatin landscape underlies key physiological changes linked to the function of particular cell types, including muscle cells and stem cells (9,10). It is also evident that disruptions to gene expression programs and chromatin modifications contribute substantially to the pathology of multiple diseases linked to hypoxia in the local environment, such as cancer, cardiovascular disease, and neurological disorders (11–13). Therefore, an improved molecular understanding of chromatin regulatory proteins and chromatin changes governing physiological responses to hypoxia is critical to further understand disease and potential therapeutic interventions.

The budding yeast *Saccharomyces cerevisiae* are facultative anaerobes, able to switch to fermentation in the absence of oxygen to generate ATP and maintain critical cell functions. Budding yeast have therefore evolved sensitive regulatory systems which allow them to maintain growth in the presence of fluctuating oxygen concentrations (14,15). This is accomplished, at least in part, through substantial changes to the transcriptome in hypoxic or anerobic conditions compared to aerobic growth (16). Key transcription factors that regulate oxygen-sensitive transcription programs have been well-characterized, including Hap1/Rox1/Mot3 (17–19), Upc2/Ecm22 (20), and Hog1 (6), among others which typically target smaller subsets of genes. Each of these transcription factors controls a unique gene expression program, although their function is also often critical to the response to other types of cellular stress. Under low or no oxygen conditions, there are multiple types of regulatory mechanisms that control the abundance and localization of these transcription factors. For example, the *HAP1* and *UPC2* mRNAs are induced relatively early in hypoxia, with *HAP1* accumulating within the first 30 minutes of hypoxic exposure and *UPC2* shortly after at 30 minutes (16,21,22). *ROX1* and *MOT3* mRNAs are reduced within 10 minutes of hypoxic exposure, consistent with their known role as repressors of hypoxia response genes (19). At the post-transcriptional level, the Hog1, Upc2, and Ecm22 proteins are all known to translocate to the nucleus in hypoxia, where they regulate gene expression. Altogether, this highlights the multiple regulatory mechanisms that act on transcription factors to ensure the proper gene expression response to limiting oxygen.

In concert with this regulation of transcription factors, chromatin modifications are also exquisitely sensitive to fluctuating oxygen concentrations, at least in part due to their dependence on key metabolites and enzymes that are oxygen sensitive. In yeast, patterns of histone modification correlate with transcriptional control mechanisms, though how this varies between hypoxic and aerobic conditions has not been thoroughly examined. For example, we have previously observed a global reduction of H4K16ac in hypoxic conditions compared to aerobic conditions (23), likely due to the reduction in acetyl coA production (24), however the extent to which histone modifications and modifying enzymes change in hypoxia in yeast to adapt to these different conditions is not known. Here, we investigated the requirement for chromatin regulators in the response to hypoxia in yeast and characterized changes to the distribution of key histone modifications in hypoxic growth, including H3K4me3 and H3K9ac marks. We identified a relatively small subset of chromatin regulators that support growth in hypoxic conditions, although there are substantial changes to histone modifications under these conditions. In particular, H3K9ac is globally reduced in hypoxia though maintains its distribution at genes dependent on their transcriptional regulation in hypoxia, whereas H3K4me3 is more abundant under hypoxic conditions and its distribution shifts from promoters to coding regions largely independent of the transcription status of genes in hypoxia. Altogether, our results provide new insights into the requirement for chromatin regulators in the yeast response to hypoxia and are the first to characterize at the genome-wide level key histone marks important to transcriptional responses to environmental changes in yeast.

## MATERIALS AND METHODS

### Yeast strains and growth conditions

All Saccharomyces cerevisiae strains used in this study are provided in Supplemental Table S1. Epitope tag integrations were made by transforming a PCR cassette amplified from the pFA6a vector (25). Double mutant strains were generated by haploid mating, sporulation and tetrad dissection. Genotypes were confirmed by growth on selective media and colony PCR. Yeast cells were grown in standard rich media (YPD: 1% yeast extract, 2% peptone, 2% dextrose).

For spot assays, yeast cells were grown to saturation overnight shaking at 30°C, then diluted next day to ∼0.2 OD_600_ to mid-log phase (∼0.4 – 0.6 OD_600_). 0.1 OD_600_ of cells was then serially diluted onto YPD plates and grown at 30°C for 2 days aerobic conditions. For hypoxic conditions, the plates were incubated at 30°C for 7 days in vacuum-sealed pouches with Wallaby oxygen absorbers (400cc) (∼ 1% O2).

### Immunoblotting

Strains were grown overnight to saturation in 3 mL of YPD, then diluted to ∼0.2 OD_600_ and grown to mid-log phase (∼0.4 – 0.6 OD_600_) for aerobic conditions. For hypoxic conditions, the overnight culture was diluted and grown to mid-log phase in YPD in hypoxic pouches for 18 hours. 2.5 OD_600_ units of culture were spun down at 10,000 rpm for 2 minutes, and the pellet was resuspended in 100 µL of H_2_O and 100 µL of 0.2 M NaOH. Cells were incubated with NaOH for 5 minutes at room temperature. Cells were then collected with centrifugation at 10,000 rpm for 2 minutes, and the pellet was resuspended in 50 µL of 1X SDS loading buffer, then boiled for 5 minutes. Before loading for SDS PAGE, cell debris was pelleted by centrifugation.

Samples were loaded on SDS-PAGE and proteins were transferred onto Immobilon-FL polyvinylidene difluoride membrane (Sigma, Cat #IPFL00005). Membranes were probed with the appropriate antibody overnight and blots were visualized using the Odyssey CLx Imager. Total protein stain was performed with Revert 700 total protein stain kit (LICOR, Cat #926-11010). The following antibodies were used for immunoblotting: mouse anti-FLAG (Sigma, Cat #F1804), mouse anti-c-Myc (Invitrogen, Cat #MA1-980), and IRDye 680RD goat anti-mouse IgG (LiCor, Cat #926-68070).

### Reverse transcriptase-quantitative PCR (RT-qPCR)

RNA was extracted as previously described (23,26,27) from log phase cultures (OD_600_∼0.4 – 0.8) grown under aerobic or hypoxic conditions (18 hrs) and at indicated time points in aerobic conditions. MasterPure Yeast RNA Purification Kit (Biosearch, Cat #MPY03100) was used to extract RNA per manufacturer’s instructions. Genomic DNA was digested using TURBO DNA-free Kit (Invitrogen, Cat #AM1907). The Accuris qMAX cDNA Synthesis Kit (Accuris, Cat #PR2100-C-100) was used to generate cDNA from generated 0.5 – 1 µg of RNA. Transcript levels were measured using quantitative PCR (qPCR), in which 0.5 µL of cDNA and target gene primers (Supplemental Table S2) were added to 10 µL reactions containing qMax Green Low Rox qPCR Mix (Accuris, Cat #PR2000-L-1000). Amplification was performed using a Bio-Rad CFX384 Real-time Detection System. Three biological replicates and three technical replicates per primer set were performed. Gene expression values were normalized to *TFC1*, which has stable mRNA levels under multiple growth conditions (28). Statistical significance was calculated with 2-way ANOVA. P value is indicated as follows: * ≤ 0.05, ** ≤ 0.01, *** ≤ 0.001.

### Chromatin Immunoprecipitation Sequencing (ChIP-Seq)

Chromatin immunoprecipitation was performed as described (23,29,30). Briefly, cells were grown in 200 mL of YPD at to mid-log phase 30°C either with shaking or in BD GasPak EZ anaerobe pouches. Cultures were then fixed with 1% formaldehyde (Sigma, Cat #F8775) for 20 minutes at room temperature. Formaldehyde was then quenched by adding 10 mL of 2.5 M glycine and shaking for an additional 5 minutes. Cells were lysed with 2 mL of chIP lysis buffer (50 mM Tris-HCl pH 7.5, 150 mM NaCl, 1 mM EDTA, 1% NP-40) and bead beat with 500 µL acid-washed glass beads for 5 minutes at 4°C. Chromatin was pelleted by centrifugation at 13,000 rpm for 20 min at 4°C. Chromatin was then resuspended in 250 µL MNase digestion buffer (10 mM Tris-HCl pH 7.5, 10 mM NaCl, 3 mM MgCl_2_, 1 mM CaCl_2_, 1% NP-40), and 2.5 µL of MNase (Worthington, Cat #LS004797) was added. MNase digestion occurred at 37°C for 20 minutes, and the reaction was stopped via addition of 5 µL of 0.5M EDTA. Extract was clarified via centrifugation at 13,000 rpm for 15 minutes at 4°C. Chromatin was normalized via Bradford assay to 120 µg in 1.1 mL and 100 µL was removed as the input. To the remainder of the normalized chromatin, 30 µL of protein A/G magnetic beads (Pierce, Cat #88802), pre-incubated with 5 µg of the appropriate antibody, were added and incubated overnight at 4°C with rotation. The following antibodies were used for chIP: rabbit anti-H3 (abcam, Cat #ab1791), rabbit anti-H3K9ac (Sigma, Cat #06-942), and rabbit anti-H3K4me3 (Epicypher, Cat #13-0041).

Protein-DNA complexes were eluted using 1% SDS and 0.1 M NaHCO_3_, cross-links were reversed, and samples were treated with proteinase K (RPI, Cat #P50220) and RNase A (Sigma, Cat #R-5125). DNA was purified using the MinElute PCR purification kit (Qiagen, Cat #28004). DNA concentration was measured with Qubit. Sequencing libraries were prepared using Illumina reagents at a commercial vender (Azenta) and next generation sequencing was performed on an Illumina Novaseq 6000 at Azenta.

### ChIP-seq analysis

For ChIP-Seq analysis, FASTQ files were uploaded to the Galaxy server and subsequent analysis was performed in Galaxy (31). Sequencing adaptors were trimmed using TrimGalore and reads were filtered based on a minimum mapping quality (MAPQ) of 20. Reads from input and IP samples were mapped to the *Saccharomyces cerevisiae* genome sacCer3 using HISAT2 and to the *Drosophila melanogaster* dm6 genome. Normalization factors were calculated based on spike-in chromatin mapped reads and input samples as described (32). Peak calling was performed with MACS2 using the scaling factors calculated based on spike-in normalization. For H3K9ac peaks, broad peaks were called, whereas for H3K4me3 peaks, narrow peaks were called. MACS2 did not build the shifting model for peak calling and the q-value, or minimum FDR, was set to 0.05. Peak annotation and distribution relative to genomic features was determined using ChIPseeker, where the TSS-associated region was set to −800 bp to +500 bp. Metagene plots were generated using computematrix and plotProfile or plotHeatmap. Bigwig files were generated using bamcoverage and appropriate scaling factors calculated based on spike-in and input normalization. Genome tracks were visualized using the Integrative Genomics Viewer (IGV) (33). Additional analysis was performed using Excel or GraphPad Prism.

## RESULTS

### Genome reprogramming and requirements for histone modifying complexes in response to hypoxia

Upon transition of budding yeast from aerobic to hypoxic environments, an initial, rapidly induced gene expression program provides an early response to low oxygen conditions and promotes cell survival (16,34). Following prolonged hypoxia exposure, this early response transitions to an adapted response in which metabolic programming is shifted to a fermentative state to support survival in low oxygen (16,34). We previously profiled yeast cells following 18 hours growth in hypoxia in rich media (approximately 1% oxygen) and compared them to cells continuously grown in aerobic conditions (23). In these cells adapted to hypoxic conditions, we observed massive reprogramming of the genome, including more than 1000 genes upregulated and more than 800 genes downregulated compared to aerobic conditions, representing close to one-third of the yeast genome. As expected, genes encoding factors required for energy-dependent processes such as protein translation are generally downregulated, as are genes important for progression of the cell cycle, which typically slows during hypoxic growth (**Figure 1A**). Genes that are upregulated in hypoxic conditions include those associated with transport across membranes, metabolism of lipids, carbohydrates, and amino acids, and organization of the cell wall, which shows altered composition in low oxygen conditions. Additionally, genes required for respiration and energy maintenance under hypoxic conditions and genes associated with the response to starvation are upregulated (**Figure 1A**; (23)). Altogether, the magnitude of the changes in gene expression observed in cells adapted to hypoxia underscores the massive metabolic reprogramming that occurs in these conditions.

**Figure 1.**
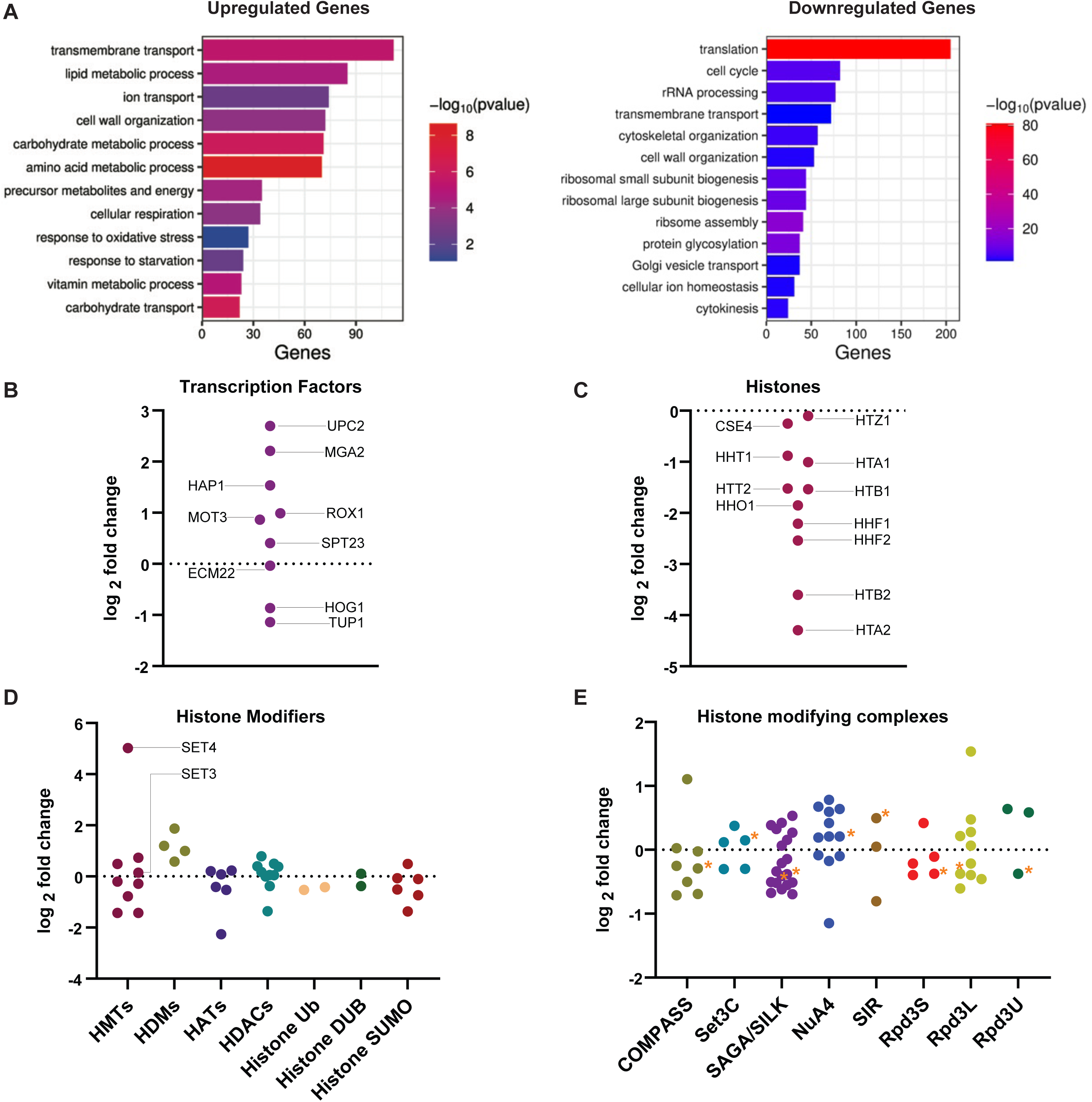
Genomic reprogramming during hypoxia impacts expression of transcription factors and chromatin regulators. **A.** Gene ontology of differentially expressed genes in *WT* cells grown in hypoxic conditions compared to aerobic conditions from RNA-Seq (23). Gene counts for each category and −log_10_ of p-values are shown. Graphs of the log_2_ fold change of transcription factors known to regulate hypoxic gene expression (**B**) and histones (**C**) from RNA-seq of *WT* cells under hypoxic conditions. **D.** Graph of log_2_ fold change of families of histone-modifying proteins: HMT (histone methyltransferase), HDM (histone demethylase), HAT (histone acetylase), HDAC (histone deacetylase), Ub (ubiquitin ligase), DUB (deubiquitinase), and SUMO (SUMO ligases and proteases). **E.** Graph of log_2_ fold change of complex members associated with some of the histone-modifying proteins in (**D**). Asterisk denotes log_2_ fold change value of the catalytic subunit of each complex.

We next assessed the expression of key transcription factors and chromatin modifiers in hypoxic conditions to better define the role for these regulators in adaptation to hypoxia. In our RNA-seq analysis, we observed that the major transcription factors that activate the hypoxic gene expression program in response to depleted sterols and heme show some modest changes in gene expression. Primarily, we detected upregulation of the sterol response factor Upc2, although the related transcription factor Ecm22 does not show any change in abundance (**Figure 1B**). Another key regulator of lipid metabolism, Mga2, which translocates to the nucleus in hypoxia (35), shows substantial upregulation under these conditions, as does the heme responsive transcription factor Hap1. Interestingly, although Mot3, Rox1, and Tup1 are all implicated in repression of hypoxia-induced genes, only the co-repressor Tup1 is downregulated at the level of mRNA under these conditions.

Next, we investigated patterns of expression for genes encoding histones and the major chromatin-modifying families in yeast such as histone acetyltransferases (HATs), histone deacetylases (HDACs), histone methyltransferases (KMTs), and histone demethylases (KDMs), among other key modifiers. We first noted that all the histone genes are downregulated in hypoxic conditions (**Figure 1C**), which is most likely a consequence of the slowed cell cycle under hypoxia and therefore a reduced requirement for new histone synthesis (15,16,34). For the majority of chromatin modifiers evaluated, there was not a unified expression pattern when comparing hypoxic conditions to aerobic conditions, with the exception of the KDMs. These enzymes are all upregulated in hypoxia, which most likely represents a compensatory mechanism that may counteract their reduced catalytic activity under limiting oxygen, as has been demonstrated for some KDMs in mammalian systems (7,36–38). Otherwise, the expression of the majority of histone-modifying enzymes changes modestly in long-term hypoxic conditions, with most of them showing no change or downregulation, including the most downregulated of the HATs, *RTT109*, and HDACs, *HST3*. The most upregulated of all of the chromatin-related genes evaluated is the SET domain-containing protein *SET4*, which is uniquely expressed in hypoxia and is known to regulate hypoxia response genes (23,39). Furthermore, we evaluated the expression of non-enzymatic components of multiple chromatin-modifying complexes to determine whether any complexes are coordinately regulated in hypoxia-adapted cells (**Figure 1E**). Here, we did not observe a coordinated change in expression for any particular complex, and found that the most upregulated non-enzymatic component of the complexes evaluated is *CTI6*, a component of the Rpd3L complex.

To define the requirement for these regulatory transcription factors and chromatin modifiers in the adaptation to hypoxia, we performed growth assays on plates incubated in either aerobic conditions or hypoxic conditions for up to 7 days. We first tested deletions of many of the major transcription factors required for regulating the hypoxic gene expression response and observed only very mild impacts on growth in hypoxic conditions compared to wildtype (**Figure 2A**), with the exception of loss of *TUP1*, which shows improved growth in hypoxia potentially due to the additional de-repression of hypoxia response genes in this mutant (40). Next, we evaluated the impact of the loss of the non-essential components of several major chromatin regulatory families, including the KMTs (**Figure 2B**), KDMs (**Figure 2C**), HATs (**Figure 2D**), and HDACs (**Figure 2E**). We found that the majority of the deletion mutants for these chromatin factors showed little change in growth compared to wildtype under hypoxic conditions. There were a small number of factors that do appear to contribute to cell survival or growth in hypoxia, which includes the SET domain containing proteins Set3 and Set4 (**Figure 2B**), the HATs Gcn5 and Rtt109 (**Figure 2D**), and the HDACs Sir2 and Rpd3 (**Figure 2E**), with *rpd3Δ* cells showing the most severe growth defect under hypoxic conditions. These observations are largely consistent with known roles for a number of these factors, including Set4, Gcn5/SAGA, and Rpd3 in both activation and repression of genes regulated based on oxygen availability (1)(6,41).

**Figure 2.**
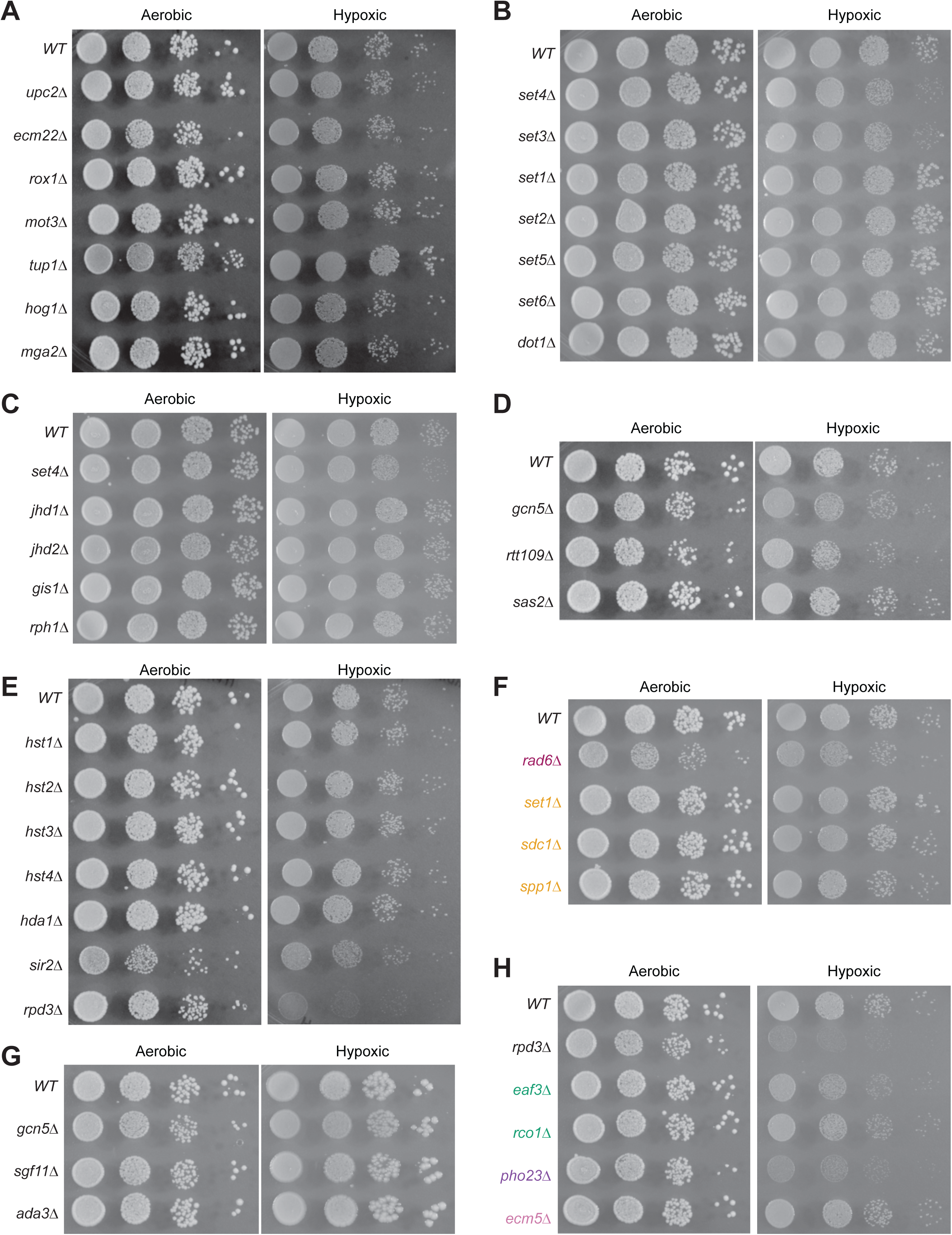
Growth under hypoxia depends on a select few histone modifiers and transcriptional regulators. Spot assays of 10-fold serially diluted cells grown on YPD plates at 30°C at either aerobic (2 days) or hypoxic (7 days) conditions, grouped based on the following categories: (**A**) transcription factors, (**B**) histone methyltransferases, (**C**) histone demethylases, (**D**) histone acetyltransferases, (**E**) histone deacetylases, (**F**) proteins affecting H3K4 methylation (maroon denotes H2B ubiquitin machinery, orange denotes COMPASS complex), (**G**) SAGA complex members, and (**H**) Rpd3 complex members (green denotes Rpd3S, purple denotes Rpd3L, and pink denotes Rpd3µ).

Given that we observed differential regulation of expression of some chromatin complex components in hypoxia (**Figure 1E**), we also evaluated whether individual components of some complexes show a different requirement for growth in hypoxia. For example, we tested deletions of components of the COMPASS complex which differentially regulate H3K4 methylation status, as well as a *rad6Δ* mutant, which does not have H2B ubiquitination, inhibiting H3K4me deposition. Of these strains, we observed a mild defect in growth in an *spp1Δ* mutant strain (**Figure 2F**) which specifically lacks H3K4me3 and has roles in DNA replication and repair (42); however there was no clear growth defect specifically in hypoxia in the other COMPASS mutants tested. We also tested additional deletion mutants of the SAGA complex (**Figure 2G**), including *sgf11Δ*, which disrupts the deubiquitination activity of SAGA, and *ada3Δ*, another component of the HAT module, but did not observe any substantial impact on growth of these cells in hypoxia, despite the known role for SAGA in regulating hypoxia response genes and stress response genes in general (43). The HDAC Rpd3 exists in multiple complexes in yeast (44), with the Rpd3L complex largely responsible for regulating transient and inducible stress responses (45) and the Rpd3S complex acting as a repressor of cryptic transcription, among other processes (46). We evaluated deletion mutants of several components of these complexes and determined that a *pho23Δ* mutant showed the strongest defect in growth in hypoxia, similar to the *rpd3Δ* cells (**Figure 2H**). Pho23 is a core component of the Rpd3L complex (47,48), indicating that the requirement for Rpd3 in the response to hypoxia is largely driven by the functions of the Rpd3L complex rather than other Rpd3-containing complexes.

### The Set4 and Set3 paralogs are differentially regulated in hypoxia and independently contribute to survival and gene regulation in hypoxia

While Set4 has been previously reported to be important for cell survival and gene regulation in hypoxia (23,39), our screen of transcription and chromatin mutants in hypoxia showed that loss of its paralog, Set3, causes a similar growth defect in hypoxia (**Figure 2B**). *SET4* is highly upregulated in hypoxic conditions (**Figure 1D**, (23,39)) and the protein is largely only detectable under hypoxic conditions (**Figure 3A**). Conversely, Set3 does not show any oxygen concentration-dependent regulation at the level of mRNA or protein abundance (**Figure 3A**). In addition, as cells transition from hypoxic conditions back to aerobic conditions, the *SET4* mRNA is quickly downregulated (**Figure 3B**) and previous work has shown that the Set4 protein levels also decrease rapidly (39), indicating that cells specifically require Set4 in conditions of low oxygen. To test whether Set4 and Set3 have similar functions in hypoxic conditions, we generated *set3Δ set4Δ* double mutant cells and tested their growth under aerobic and hypoxic conditions. While these double mutants grew indistinguishably from wildtype under normal, aerobic conditions, they showed an enhanced growth defect in hypoxia compared to either single *set3Δ* or *set4Δ* mutants (**Figure 3C**). Furthermore, we assessed gene expression of hypoxia response genes, particularly those associated with cell wall maintenance that are known targets of Set4 (23), and found that while some genes such as *ECM13* and *SED1* show dependence on both Set4 and Set3 for their expression, many others are regulated uniquely by Set4, but not Set3, including *ERG4*, *DAN4*, and *CWP2* (**Figure 3D**). Altogether, these data suggest that while Set4 and Set3 may share some overlapping gene targets during growth in hypoxia, their molecular functions are distinct and they are each independently required for cell survival in hypoxic conditions.

**Figure 3.**
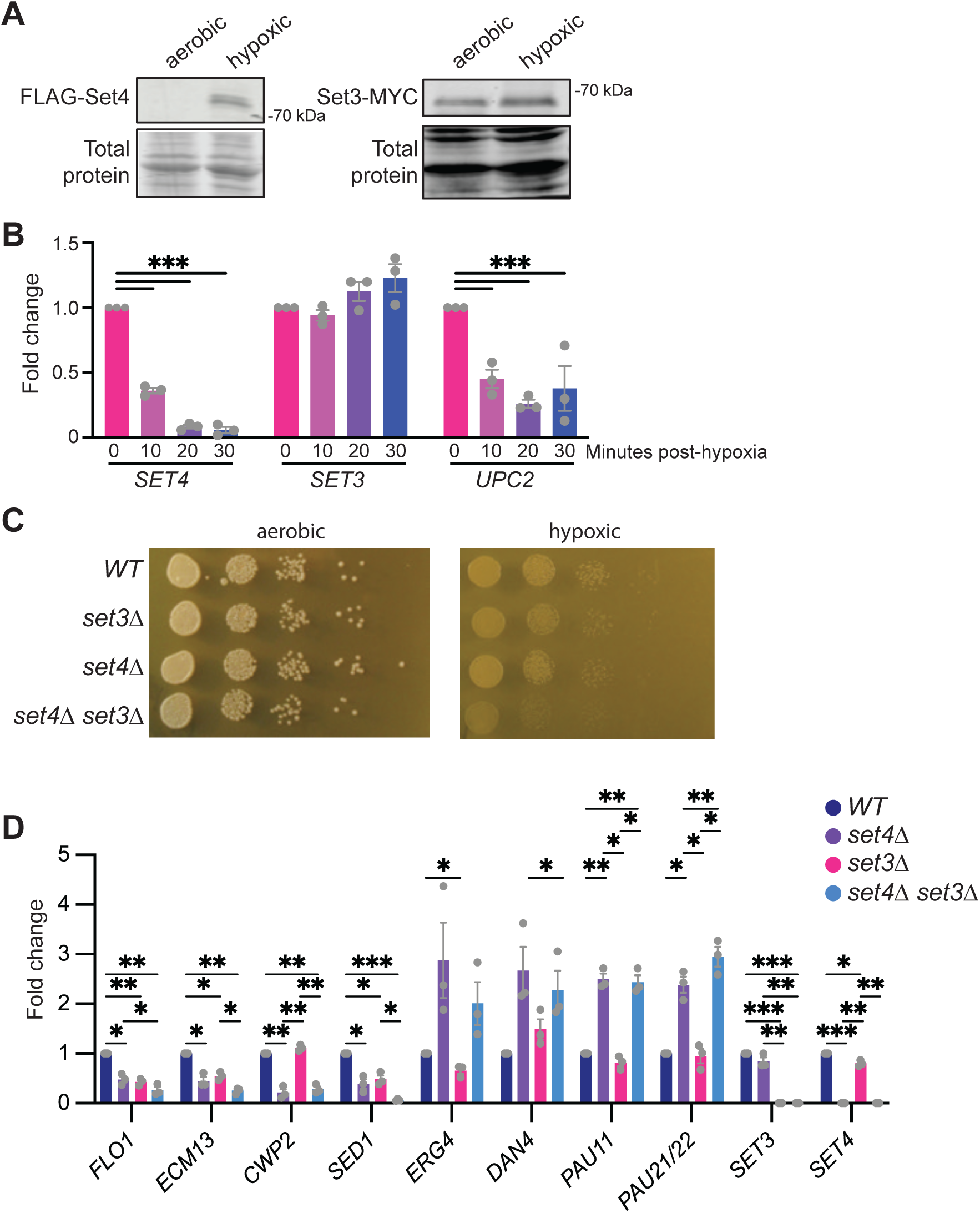
Set4 and its paralog Set3 play unique roles in hypoxia. **A.** Immunoblots depicting FLAG-Set4 (left), Set3-MYC (right), and total protein stain from cells grown under aerobic or hypoxic conditions. **B.** qRT-PCR showing mRNA expression SET4, SET3, and UPC2 of WT cells grown under hypoxic conditions for 18 hr and then exposed to aerobic conditions for the indicated time points. Fold change normalized to *TFC1* levels. **C.** Spot assay of 10-fold serial dilutions of *WT*, *set3Δ*, *set4Δ*, and *set4Δ set3Δ* strains grown on YPD plates grown for 2 days under aerobic conditions or 7 days under hypoxic conditions at 30°C. **D.** qRT-PCR showing mRNA expression of cell wall genes in *WT*, *set3Δ*, *set4Δ*, and *set4Δset3Δ* strains grown under aerobic or hypoxic conditions. Fold change normalized to TFC1 levels. Statistical significance was calculated with 2-way ANOVA. P value is indicated as follows: * ≤ 0.05, ** ≤ 0.01, *** ≤ 0.001.

### Hypoxia induces genome-wide changes to the abundance and distribution of H3K9ac and H3K4me3 modifications

Our evaluation of the requirement for chromatin regulators in hypoxia revealed that a relatively small subset acts uniquely to maintain cell survival under hypoxic conditions, such as Set4, Set3, and Rpd3. However, the metabolic shift induced under hypoxic conditions impacts some histone-modifying enzymes directly, including demethylases, and alters the pool of metabolites required for modifications, particularly acetyl-coA (49). Due to the large number of regulatory and metabolic changes, the chromatin landscape has been shown to be substantially altered in cells exposed to hypoxia in different mammalian cell types, and especially in cancer cell lines from solid tumors (12,50–52). However, the impact on chromatin modifications in yeast adapted to hypoxic growth has not been reported. To better define this chromatin state, we profiled two major histone modifications by chIP-sequencing, H3K4me3 and H3K9ac, and used both spike-in and input-based normalization (32) to define changes to the abundance and distribution of these marks genome-wide under hypoxic conditions relative to aerobic conditions.

In wildtype yeast grown at approximately 1% O_2_ for 18 hours, we observed decreased immunoprecipitation of chromatin using an antibody against H3K9ac from cells compared to cells grown in aerobic conditions (**Figure 4A**) and metagene analysis of H3K9ac across all genes shows substantially less H3K9ac across promoters and gene bodies during hypoxia compared to aerobic growth, although still maintaining its expected enrichment near transcription start sites (TSSs) in both conditions (**Figure 4B**). The reduced abundance of H3K9ac is also evident in normalized genome browser tracks (**Figure 4C**), which show decreased H3K9ac throughout the region, particularly at genes that are downregulated in hypoxic conditions, such as *YOL057W* and *GPM3*. Genes in this region which are upregulated during hypoxia, such as *ARG1* and *DDR2*, show less change in the pattern of H3K9ac occupancy. In comparing genes containing H3K9ac peaks in aerobic and hypoxic conditions, we observed a high proportion of shared peaks; however there were more peaks identified under hypoxic than aerobic conditions (**Figure 4D**). The distribution of these peaks relative to genomic elements is largely similar, with the majority localizing to promoters near TSSs under both conditions, with some occupancy in gene bodies (primarily exons) that moderately increases under hypoxic conditions (**Figure 4D**). We evaluated the distribution of H3K9ac relative to genes identified by RNA-seq to be differentially expressed in hypoxic conditions (**Figure 1A**, (23)). We found that new peaks of H3K9ac in hypoxia were associated with genes both up-and down-regulated in hypoxia (**Figure 4E**). These peaks largely corresponded to the major categories of genes that are differentially expressed in hypoxia, including downregulated genes linked to translation and the cell cycle, and upregulated genes associated with amino acid, carbohydrate, and lipid metabolism, intracellular transport, and cellular respiration (**Figure 4E**). However, there are also a large number of peaks of H3K9ac gained in hypoxia that are not associated with a significant change in gene expression of neighboring genes at the time point monitored (**Figure 4E**), though these genes do not show enrichment for any particular biological processes or function. These data demonstrate the global decrease in H3K9ac in hypoxia and suggest its change in abundance is not clearly linked to gene expression under these conditions in yeast.

**Figure 4.**
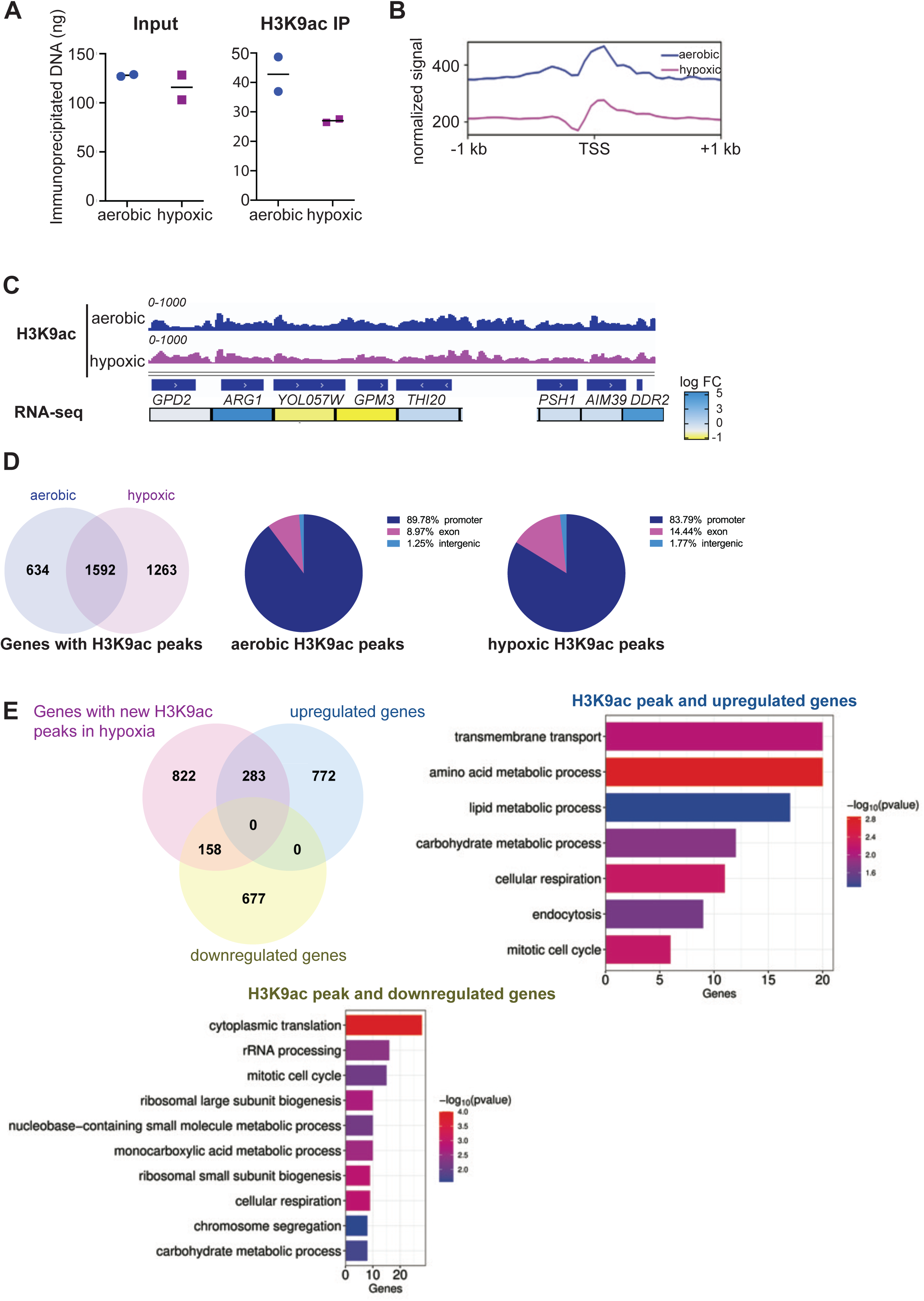
H3K9ac is less abundant throughout the genome in hypoxia. **A.** Comparison of DNA obtained after immunoprecipitation with H3K9ac antibody in aerobic and hypoxic conditions, along with the amount of DNA present in input controls (n = 2 aerobic, n = 2 hypoxic biological replicates). DNA concentration was measured using Qubit. **B.** Metagene plot of average H3K9ac ChIP-Seq signal at TSSs ± 1 kb at all genes. **C.** ChIP-Seq genome browser tracks of H3K9ac mapped reads and RNA-Seq log2 fold-change at promoter and intergeneic regions (bottom). **D.** Venn diagram showing overlap of genes containing H3K9ac peaks in either aerobic or hypoxic conditions (left) and percentage distribution of aerobic or hypoxic H3K9ac peaks of genomic features (middle and right). **E.** Venn diagram showing overlap of genes that gained new H3K9ac peaks under hypoxic conditions with genes that were either upregulated or downregulated in the RNA-Seq (left). Gene ontology analysis of overlapping 283 upregulated genes (right) or overlapping 158 downregulated genes (bottom).

We next analyzed the distribution of another histone modification linked to regulation of environmentally responsive genes, H3K4me3. In wildtype yeast, immunoprecipitation of chromatin using anti-H3K4me3 antibody showed significantly more DNA bound to H3K4me3 under hypoxic conditions than aerobic conditions (**Figure 5A**). In both aerobic and hypoxic conditions, H3K4me3 was identified largely at transcription start sites (TSSs), however, across all genes, there was substantially more H3K4me3 bound in hypoxic conditions than aerobic conditions and it appeared to spread more into gene bodies in hypoxia (**Figure 5B**). Inspection of specific genomic regions, including genes that are induced or repressed in hypoxia, generally showed increased abundance of H3K4me3 and further spreading of the mark away from the TSS into coding sequences (**Figure 5C**). Indeed, identification of genomic regions bound by H3K4me3 showed predominantly promoter localization under both aerobic and hypoxic conditions. However, there was a substantial increase in H3K4me3-bound exons or coding sequences in hypoxia (**Figure 5D**). Gene ontology analysis did not reveal any functional categories of genes that were enriched in the population of genes carrying H3K4me3 in coding sequences. Consistent with the increased exon-bound H3K4me3 in hypoxia, the distance of H3K4me3 peaks from the TSS in hypoxic-grown cells increases, with more enrichment of peaks downstream of the TSS into coding sequences (**Figure 5E**), indicative of spreading of this mark under limiting oxygen, as observed in other systems (37,52,53).

**Figure 5.**
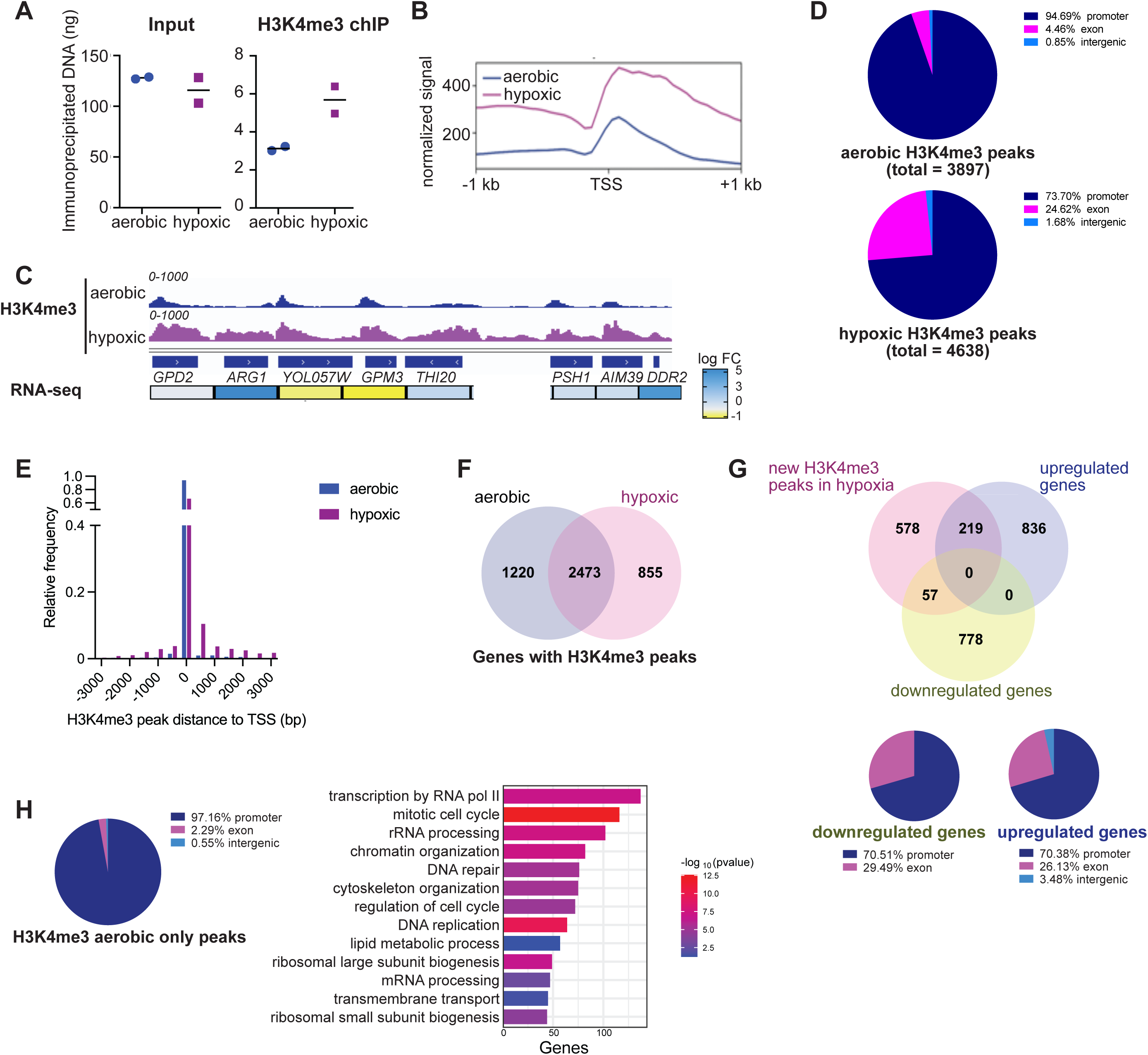
H3K4me3 increases in abundance in hypoxia. **A.** Comparison of DNA obtained after immunoprecipitation with H3K4me3 antibody in aerobic and hypoxic conditions, along with the amount of DNA present in input controls (n = 2 aerobic, n = 2 hypoxic biological replicates). DNA concentration was measured using Qubit. **B.** Metagene plot of average H3K4me3 ChIP-Seq signal at TSSs ± 1 kb at all genes. **C.** ChIP-Seq genome browser tracks of H3K4me3 mapped reads (top) and RNA-Seq log2 fold-change at promoter and intergeneic regions (bottom). **D.** Distribution of H3K4me3 peaks under aerobic (top) or hypoxic conditions (bottom) across genomic features. **E.** Relative frequency of distance of H3K4me3 peaks relative to TSS in aerobic or hypoxic conditions. **F.** Venn diagram showing overlap of genes containing H3K4me3 peaks across aerobic and hypoxic conditions. **G.** Venn diagram showing overlap of genes that gained new H3K4me3 peaks under hypoxic conditions with upregulated and downregulated genes in hypoxia obtained from RNA-Seq data (top) and the distribution of H3K4me3 peaks across genomic features in the overlapping 57 downregulated genes or 219 upregulated genes (bottom). **H.** Distribution of the 1220 H3K4me3 peaks that were found in aerobic but not in hypoxic conditions across genomic features (left) and gene ontology analysis of the genes associated with those peaks.

A comparison of genes with associated H3K4me3 peaks showed a substantial fraction of genes are shared between the aerobic and hypoxic conditions; however, there are unique peaks in both conditions, with more peaks identified under aerobic conditions despite the lower abundance of the mark overall (**Figure 5F**). The genes which gained H3K4me3 peaks in hypoxia are more highly associated with genes upregulated in hypoxic conditions compared to those downregulated (**Figure 5G**), however they both share a largely similar distribution relative to the TSS with a high percentage of promoter-bound H3K4me3 but also more than a quarter of genes with H3K4me3 in coding sequences. Interestingly, gene ontology analysis did not reveal enrichment of particular functional categories of the genes gaining H3K4me3 in hypoxia that were identified as differentially expressed at this timepoint in hypoxia (**Figure 1A**, (23)). However, we did observe that the H3K4me3 peaks unique to aerobic conditions and lost in hypoxia were predominantly associated with promoter-bound H3K4me3 and genes typically downregulated in hypoxia, including those involved in cell cycle regulation, transcription, ribosome biogenesis, and lipid metabolism (**Figure 5H**). Altogether, these data highlight the undirected gain of H3K4me3 throughout the genome in hypoxic conditions and its decreased specificity for its stereotypical TSS-associated binding.

## DISCUSSION

It is well established in eukaryotic cell types that limited oxygen availability induces substantial changes to gene expression programs via the activity of oxygen-responsive transcription factors to drive programs that facilitate cell survival and adaptability. In addition, the chromatin landscape is highly sensitive to changes in oxygen availability through impacts on signaling pathways, direct sensing of oxygen by chromatin-modifying enzymes, primarily demethylases, and through regulation of the availability of enzymatic co-factors, such as acetyl-coA (49). However, our understanding of the altered histone modification status and the function of chromatin regulators in hypoxic conditions, particularly on a genomic scale, is still limited. While chromatin modification status in hypoxia has been described in numerous disease models including cancer cell lines, tumors, and models of cardiovascular disease (5,12,37,50,53), the genome-wide impact of hypoxia on chromatin in model systems, such as the facultative anaerobe budding yeast, has not been thoroughly explored. Here, we presented new insights into the regulation and growth requirement for key chromatin modifiers in hypoxia in yeast and describe the genomic changes to H3K9ac and H3K4me3 under these conditions. This work also serves as a foundation for further investigation of roles and functional properties of conserved chromatin modifiers in low oxygen cellular environments.

In assessing transcriptional regulation of the major hypoxia-responsive transcription factors and chromatin-modifying enzymes and complexes in yeast at steady state during hypoxia (16), we did not observe much change in mRNA abundance for the majority of factors evaluated (**Figure 1**). Furthermore, when we analyzed expression levels of different components of major chromatin regulatory complexes, there was little uniformity in the expression patterns of individual complex components. Previous work indicates that several of these transcription factors, such as Upc2, Ecm22, Mga2, and Hap1, are post-translationally regulated in response to environmental shifts (21,22,35,54), which may also extend to some of the chromatin modifiers (55). However, this suggests that while cellular hypoxia responses are driven in large part through regulation of transcription programs, this does not encompass the transcription factors and chromatin regulators responsible for controlling this program, though there may be dynamic changes in expression for these factors at different timepoints in the hypoxic response. One clear exception to this observation is the family of JmjC-domain containing histone demethylases, which depend on oxygen for their catalytic activity, and are all uniformly upregulated in hypoxia in yeast. Similar upregulation of histone demethylases in other systems has been postulated to compensate for reduced catalytic activity under low oxygen conditions (12,36,38,49), however the extent to which this potential compensatory mechanism is employed and its effectiveness at counteracting changes to histone methylation abundance has not been fully explored.

Beyond the histone demethylases, the SET domain containing protein Set4 is highly upregulated at the transcriptional level in hypoxia, as previously described (23,39). We also demonstrate that the mRNA encoding *SET4* is rapidly turned over if cells are shifted from hypoxic back to aerobic conditions (**Figure 3**). This suggests unique regulation of the *SET4* gene in hypoxia and highlights its specific function under these conditions, which distinguishes it from many other general chromatin modifiers and from its paralog, Set3 (56). Interestingly, while *SET3* does not appear to be regulated at the transcriptional level during hypoxia, loss of Set3 does disrupt cell growth and gene expression in hypoxia and cells lacking both Set3 and Set4 have an enhanced growth defect under hypoxic conditions, indicating that they each contribute to survivability in hypoxia. We previously observed a similar genetic interaction between *set3*Δ and *set4*Δ mutants in oxidative stress (57,58). Our analysis of gene expression in single and double *SET3* and *SET4* mutants indicates that they do share some target genes, though they also distinctly regulate other genes, indicating that they are acting via different molecular mechanisms. Further investigation of the functions of both Set3 and Set4 in hypoxia will distinguish the contributions of their shared and unique roles to hypoxic gene regulation.

To better understand the chromatin landscape in yeast during hypoxia, we evaluated the distribution of two key chromatin marks in yeast that are typically linked to genomic responses to different types of stress, H3K9ac and H3K4me3. H3K9ac is regulated by the SAGA histone acetyltransferase complex and the Rpd3 histone deacetylase, which typically work coordinately to establish H3K9ac distribution particularly as gene expression programs change during stress (6,41,43–45,59). We did observe that yeast specifically rely on the function of the Rpd3L complex for survival in hypoxia, though there is only a modest decrease in survival in mutants of the SAGA complex (**Figure 2**). While we previously observed reduced bulk levels of histone acetylation in hypoxia (23), as expected due to the reduced availability of acetyl coA, this change in acetylation abundance had not been mapped genome-wide in yeast. Our data show here a global reduction in H3K9ac, though it generally maintains a similar distribution pattern under hypoxic conditions as under aerobic conditions. H3K9ac is enriched near TSSs, though there is also a broader distribution into gene bodies. Interestingly, more unique peaks are identified under hypoxic conditions than aerobic conditions, potentially due to an overall reduction in broad domains containing H3K9ac. However, genes gaining H3K9ac under hypoxic conditions were both up-and downregulated relative to aerobic conditions at the timepoint tested, and there was no particular enrichment for functional gene categories at other regions gaining H3K9ac under hypoxia. This suggests that while there is an overall decrease in H3K9ac throughout the genome, it may be generally nonspecific and it does not appear directly linked to gene expression under these conditions.

Conversely, H3K4me3 is more abundant at chromatin under hypoxic conditions compared to aerobic conditions in yeast. A similar increase in histone methylation has been reported in other organisms and disease models (7,8,53), and has been attributed to the reduction in catalytic activity of demethylases in hypoxia. Jhd2 is the demethylase for H3K4me3 in yeast, and while its catalytic activity is dependent on oxygen, the extent to which it is inhibited in yeast grown in hypoxia is not known. However, the distribution of H3K4me3 observed here in hypoxia shows similar alterations to the H3K4me3 distribution in the absence of Jhd2 (60,61), with increased spreading away from the TSS and into coding sequences. Therefore, it is expected that this H3K4me3 pattern is established under hypoxic conditions through the biochemical loss of function of Jhd2. Under these conditions, cells are able to mount widespread changes to the gene expression program, despite changes to key signals to major transcriptional regulators such as reduced promoter histone acetylation and non-specific deposition of H3K4me3 outside of the TSS. This indicates that transcriptional regulation is largely intact despite the altered chromatin landscape, though there are likely to be condition-specific alterations to the functions and mechanisms used by major chromatin-modifying complexes and key transcriptional regulators in hypoxia. Further investigation of chromatin modifying complexes and key regulators under low oxygen conditions is required to identify new or alternative mechanisms used by these factors to maintain transcriptional responses under this challenging cellular stress.

## LIST OF ABBREVIATIONS

PTM: post-translational modification
H4K16ac: histone 4 lysine 16 acetylation
H3K4me3: histone 3 lysine 4 tri-methylation
H3K9ac: histone 3 lysine 9 acetylation
PCR: polymerase chain reaction
RT-qPCR: reverse transcription-polymerase chain reaction
cDNA: copy DNA
chIP: chromatin immnoprecipitation
HDAC: histone deacetylase
KMT: lysine methyltransferase
KDM: lysine demethylase
HAT: histone acetytransferase
SET: Su(Var)3-9, Enhancer of zeste, and Trithorax
SAGA: Spt-Ada-Gcn5 acetyltransferase
COMPASS: Complex of Proteins Associated with Set1
TSS: transcription start site
Ub: ubiquitin
DUB: deubiquitinase
SUMO: small ubiquitin-related modifie

## DECLARATIONS

### Ethics approval and consent to participate

Not applicable.

### Authors’ contributions

WS and MYN contributed equally to the work and performed the majority of the experiments including the yeast genetics and growth assays, chIP-sequencing, and computational analysis. CA, VM, and JJS performed additional experiments. EMG conceptualized the work with WS and MYN, supervised the work, and wrote the manuscript with WS.

## Acknowledgements

The authors acknowledge other members of the Green lab for critical feedback on the work and manuscript. The authors also acknowledge Amol Shetty and Luke Tallon at Maryland Genomics and the University of Maryland School of Medicine for computational support.

## SUPPLEMENTARY MATERIAL

**Supplementary Table S1.** Yeast strains used in this study.

**Supplementary Table S2.** Oligos used in this study.

