## Supplemental Tables for "Alterations to the yeast chromatin landscape during hypoxia"

**Supplemental Table 1. *S. cerevisiae* strains used in this study.**

| Strain | Background | Mat | Genotype | Source |
|---|---|---|---|---|
| yEG001 | BY4741 | a | *his3Δ1 leu2Δ0 met15Δ0 ura3Δ0* | YKO |
| yEG322 | BY4741 | a | *set4::set4Δ::HIS3MX* | (1) |
| yEG1478 | BY4742 | alpha | *set3::set3Δ::NATMX* | This study |
| yEG218 | BY4742 | alpha | *set1::set1Δ::KANMX* | This study |
| yEG1420 | BY4742 | alpha | *set2::set2Δ::KANMX* | This study |
| yEG495 | BY4742 | alpha | *set5::set5Δ::KANMX* | This study |
| yEG096 | BY4741 | a | *set6::set6Δ::KANMX* | This study |
| yEG1433 | BY4742 | alpha | *dot1::dot1Δ::KANMX* | This study |
| yEG385 | BY4741 | a | *jhd1::jhd1Δ::KANMX* | This study |
| yEG383 | BY4741 | a | *jhd2::jhd2Δ::KANMX* | This study |
| yEG381 | BY4741 | a | *gis1::gis1Δ::KANMX* | This study |
| yEG387 | BY4741 | a | *rph1::rph1Δ::KANMX* | This study |
| yEG1572 | BY4741 | a | *gcn5::gcn5-HA::NATMX* | This study |
| yEG1438 | BY4742 | alpha | *rtt109::rtt109Δ::KANMX* | This study |
| yEG1649 | BY4741 | a | *sas2::sas2Δ::KANMX* | This study |
| yEG611 | BY4742 | alpha | *hst1::hst1Δ::HIS3MX* | This study |
| yEG1643 | BY4741 | a | *hst2::hst2Δ::KANMX* | This study |
| yEG1645 | BY4741 | a | *hst3::hst3Δ::KANMX* | This study |
| yEG1647 | BY4741 | a | *hst4::hst4Δ::KANMX* | This study |
| yEG1503 | BY4741 | a | *hda1::hda1Δ::NATMX* | This study |
| yEG917 | BY4742 | alpha | *sir2::sir2Δ::KANMX* | (2) |
| yEG921 | BY4742 | alpha | *rpd3::rpd3Δ :: NATMX* | (2) |
| yEG623 | BY4742 | alpha | *rad6::rad6Δ::HIS3MX* | (3) |
| yEG110 | BY4741 | a | *sdc1::sdc1Δ::KANMX* | (4) |
| yEG100 | BY4741 | a | *spp1::spp1Δ::KANMX* | (4) |
| yEG1471 | BY4742 | alpha | *sgf11::sgf11Δ::NATMX* | This study |
| yEG1540 | BY4741 | a | *ada3::ada3Δ::NATMX* | This study |
| yEG1455 | BY4741 | a | *eaf3::eaf3Δ::NATMX* | This study |

| | | | | |
|---|---|---|---|---|
| yEG1562 | BY4742 | alpha | *rco1::rco1Δ::NATMX* | This study |
| yEG1460 | BY4741 | a | *pho23::pho23Δ::NATMX* | This study |
| yEG441 | BY4741 | a | *ecm5::ecm5Δ::KANMX* | This study |
| yEG1159 | BY4741 | a | *upc2::upc2Δ::KANMX* | (2) |
| yEG1568 | BY4742 | alpha | *ecm22::ecm22Δ::NATMX* | This study |
| yEG238 | BY4741 | a | *rox1::rox1Δ::HISMX* | This study |
| yEG239 | BY4742 | alpha | *mot3::mot3Δ::KANMX* | This study |
| yEG1591 | BY4741 | a | *tup1::tup1Δ::NATMX* | This study |
| yEG1616 | BY4742 | alpha | *hog1::hog1Δ::HISMX* | This study |
| yEG1641 | BY4741 | a | *mga2::mga2Δ::KANMX* | This study |

**Supplemental Table 2. Oligos used in this study.**

| Gene | Analysis | Sequence |
|---|---|---|
| ***TFC1*** | RT-qPCR | 5’-CAGACACTCCAGGCGGTATT-3’<br>5’-ACCACGGTATTCTTTTTCCATC-3’ |
| ***DAN1*** | RT-qPCR | 5’-TCAGCAGAGACCAGCTCAAA-3’<br>5’-TTCAGCAGAACTTGCAATGG-3’ |
| ***CWP1*** | RT-qPCR | 5’-GAAGGTTCTGAGAGCGATGC-3’<br>5’-GACCCGTCCTTGATAGCGTA-3’ |
| ***UPC2*** | RT-qPCR | 5’-GTTTCGACAATGGCGCCTAT-3’<br>5’-AGAGGGCGTTGATGGAGAAA-3’ |
| ***SET3*** | RT-qPCR | 5’-GACCCACGAAACCATTACAGG-3’<br>5’-TTGCCTGATAAGCGTGCATC-3’ |
| ***SET4*** | RT-qPCR | 5’-CATTATACCGCCCCAACAAC-3’<br>5’-TTGAATTCCGTGTCATTGGA-3’ |
